# Frequency-specific suppression of locomotor components by the *white*^+^ gene in *Drosophila melanogaster* adult flies

**DOI:** 10.1101/2020.02.26.966937

**Authors:** Chengfeng Xiao, Shuang Qiu

## Abstract

The classic eye-color gene *white*^+^ (*w*^+^) in *Drosophila melanogaster* (fruitfly) has unexpected behavioral consequences. How *w*^+^ affect locomotion of adult flies is largely unknown. Here, we show that *w*^+^ selectively suppresses locomotor components at relatively high frequencies (> 0.1 Hz). The wildtype Canton-S male flies walked intermittently in circular arenas while the white-eyed *w*^1118^ flies walked continuously. Through careful control of genetic and cytoplasmic backgrounds, we found that *w*^+^ was associated with intermittent walking. *w*^+^-carrying male flies had smaller median values of path length per second (PPS) and reduced 5-min path length compared with *w*^1118^-carrying males. Additionally, flies carrying 2-4 genomic copies of mini-*white*^+^ (m*w*^+^) showed reduced median PPSs and decreased 5-min path length compared with *w*^1118^ flies, and the suppression was dependent on the copy number of m*w*^+^. Fourier transform of the time series (i.e. PPSs over time) indicated that *w*^+^/m*w*^+^ specifically suppressed the locomotor components at relatively high frequencies (> 0.1 Hz). Lastly, the downregulation of *w*^+^ in neurons but not glial cells resulted in an increased percentage of high-frequency locomotor components. We concluded that *w*^+^ suppressed the locomotion of adult flies by selectively reducing the high-frequency locomotor components.

## Introduction

Discovered in 1910, the *white*^+^ (*w*^+^) gene in *Drosophila melanogaster* (fruitfly) gives rise to dark red color in compound eyes (Morgan, 1910). The *w* mutation, which causes white color, is indeed the first identified mutation in fruitflies. Nowadays *w*^+^ is genetically modified as a marker gene called mini-*white*^+^ (m*w*^+^) and *w*^1118^, one of the mutant alleles, has been included in an isogenic line for the P element-mediated germ-line transformation (Klemenz et al., 1987; Spradling and Rubin, 1982). Tens of thousands of fly transformants have been produced by using the m*w*^+^/*w*^1118^ system. Central to the application of this system is that *w*^+^ is assumed to be exclusively responsible for eye pigmentation and there is no neural or behavioral consequence.

Only five years after the discovery of *w*^+^, Sturtevant observed that white-eyed males were discriminated against mating (Sturtevant, 1915). In 1995, Zhang and Odenwald reported that the ectopic expression of m*w*^+^ induces male-male courtship chaining (Zhang et al., 1995). Thus, *w*^+^ likely possesses extra-retinal biological functions in addition to eye pigmentation. Further studies have supported this possibility. *w*^1118^ flies have declined place memory (Sitaraman et al., 2008), reduced boundary preference in circular arenas (Xiao and Robertson, 2015), and impaired climbing ability and stress resistance (Campbell and Nash, 2001; Ferreiro et al., 2018). Compared with wildtype, female but not male *w*^1118^ flies have a shortened lifespan (Ferreiro et al., 2018; Qiu et al., 2017). With extensive genetic manipulation, it has been shown that *w*^+^ and m*w*^+^ induce fast and consistent locomotor recovery from anoxia (Xiao and Robertson, 2016), promote persistent one-way walking in circular arenas (Xiao et al., 2017b), and control male-female copulation success (Xiao et al., 2017a). These findings have emphasized that *w*^+^ is associated with at least mating behavior and locomotion. Therefore, the assumption that *w*^+^ has no neural or behavioral consequence is under questioning.

There is little evidence on how *w*^+^ modulates neural output or behavioral performance. It is intriguing that *w*^+^ and m*w*^+^ promote persistent one-way walking of adult flies in circular arenas (Xiao et al., 2017b), however, the underlying physiological basis is unclear. Mutation of *w* causes reduced content of biogenic amines in the head (Borycz et al., 2008; Sitaraman et al., 2008), but it is unknown whether these changes are related to the abnormalities of neural output or behavior. When rigidly restrained, Canton-S adult flies display episodic motor activities that can be recorded from the brains (Qiu et al., 2016). The episodic motor activities appear to be nearly rhythmic with an average periodicity of around 19 seconds, and such episodic motor activities are greatly reduced in *w*^1118^ flies (Qiu et al., 2016). These findings imply that mutation of *w* could impair the potentially rhythmic motor output from the nervous system. Several concerns, however, have yet to be addressed. How *w*^+^ is associated with rhythmic behavioral output is largely unknown. The contribution of wildtype genetic or cytoplasmic background to episodic motor activities needs to be carefully examined. In addition, the rhythmic motor activity might be inducible by physical restraint and thus the effects of *w*^+^ on behavior could be conditional.

The behavior of free-moving flies contains a large amount of locomotor information. Often the mutant flies display some forms of behavioral abnormalities that are readily observable. Analysis of such abnormalities, however, requires a deep understanding of the distribution of locomotor data, appropriate decomposition of locomotor components, and extensive computation. Behavioral consequences related to *w* mutation could be reflected as any of the four-level locomotor changes: the altered primary behavioral indices (such as location preference (Xiao and Robertson, 2015), instant speed, or walking direction (Kain et al., 2012; Xiao et al., 2017b)); modifications of secondary locomotor structures (such as path length over time, grouped locations, or orientations that are descriptive of a typical physiological feature (Qiu and Xiao, 2018)); altered locomotor dynamics over time (such as the alternation of active-inactive episodes or periodic appearance of a similar locomotor structure (Qiu et al., 2016)); and abnormal interactions with environmental cues (such as the presence of male/female partners (Xiao et al., 2017a) or salient objects).

We hypothesized that *w*^+^ selectively modified the behavioral structures of locomotion in adult flies. In the current study, we computed a locomotor parameter, the path length per second (PPS), and examined the association between PPS distribution and *w* allele. We then decomposed the time series of data (i.e. PPSs over time) into the frequency domain by Fourier transform. Through extensive genetic manipulation, we show that *w*^+^ selectively suppressed locomotor components at relatively high frequencies (> 0.1 Hz).

## Materials and Methods

### Flies

Fly strains and their sources are: Canton-S (Dr. Walker laboratory; originally BDSC #1), *w*^1118^ (Dr. Seroude laboratory), 10×UAS-IVS-mCD8∷GFP (attP2) (BDSC #32185), 10×UAS-IVS-mCD8∷GFP (attP40) (#32186), 20×UAS-IVS-mCD8∷GFP (attP2) (#32194), elav-Gal4 (#8765), repo-Gal4 (#7415), UAS-*w*-RNAi (#31088), *w*^1118^/Dp(1; Y)*B*^S^*w*^+^*y*^+^ (derived from #6891) and *w*^1118^/Dp(1; Y)*w*^+^*y*^+^ (derived from #7060) (Xiao et al., 2017b), UAS-w-eYFP (Evans et al., 2008). Fly lines generated by serial backcrossing (*w*^+^ F1-10 and *w*^1118^ F1-F10) and by introgressive hybridization (*w*^+^ F4 and *w*^1118^ F4) were described previously (Xiao and Robertson, 2015; Xiao and Robertson, 2016). All tested flies were cultured with a standard medium (cornmeal, agar, molasses and yeast) at room temperatures (21-23 °C) with 30 −70% relative humidity. A light/dark (12/12h) cycle of illumination was provided. Adult flies were collected within two days after eclosion. We used pure nitrogen gas to knockdown flies for sorting. Before testing, flies were raised in food vials for at least three additional days free of nitrogen exposure. Male flies were examined unless otherwise stated. Tested flies were 4-7 days old. Experiments were conducted during the daytime at least 3 h away from light on/off switch.

### Fly tracking

The fly tracking was performed by following a reported protocol (Xiao and Robertson, 2015). Briefly, flies were loaded into circular arenas (12.7 mm diameter 3 mm depth) with one fly per arena. Up to 128 flies could be examined at the same time. Flies were secured in a large chamber which was built for gas exposure and for video recording. A base flux of room air at 2 L/min was provided to remove the effect of a dead chamber (Bouhuys, 1964). A 5-min time period was allowed for flies to adapt to experimental settings. The locomotor activities were video-recorded and post-analyzed. We wrote scripts using OpenCV2 for fly tracking. Fly positions were calculated once per 0.2 s. For each fly, a dataset of 1500 position information from 300 s locomotion was collected. These data were used for subsequent behavioral analysis.

### Behavioral analysis

A major parameter, path length per second (PPS), was calculated based on fly positions. A simplified procedure for data processing and visualization was shown (Fig. 1). The sequential PPSs over time were treated as a time series and the Fourier transform was applied to extract the frequency information and the associated power strength. Since the frequency and power strength are reflective of a “subset” of locomotor structure, we used the term “locomotor component” to describe the unit structure that carries a specific frequency and power. In several experiments, the full spectrum data (i.e. all frequencies vs power) and linear regression lines were presented. For statistical convenience, we also sampled the top-ten frequency components that carried ten-highest power from full-spectrum data and examined the distribution of top-ten frequencies. Fourier transform, data visualization, and sampling were performed using the related R packages.

**Figure 1.**
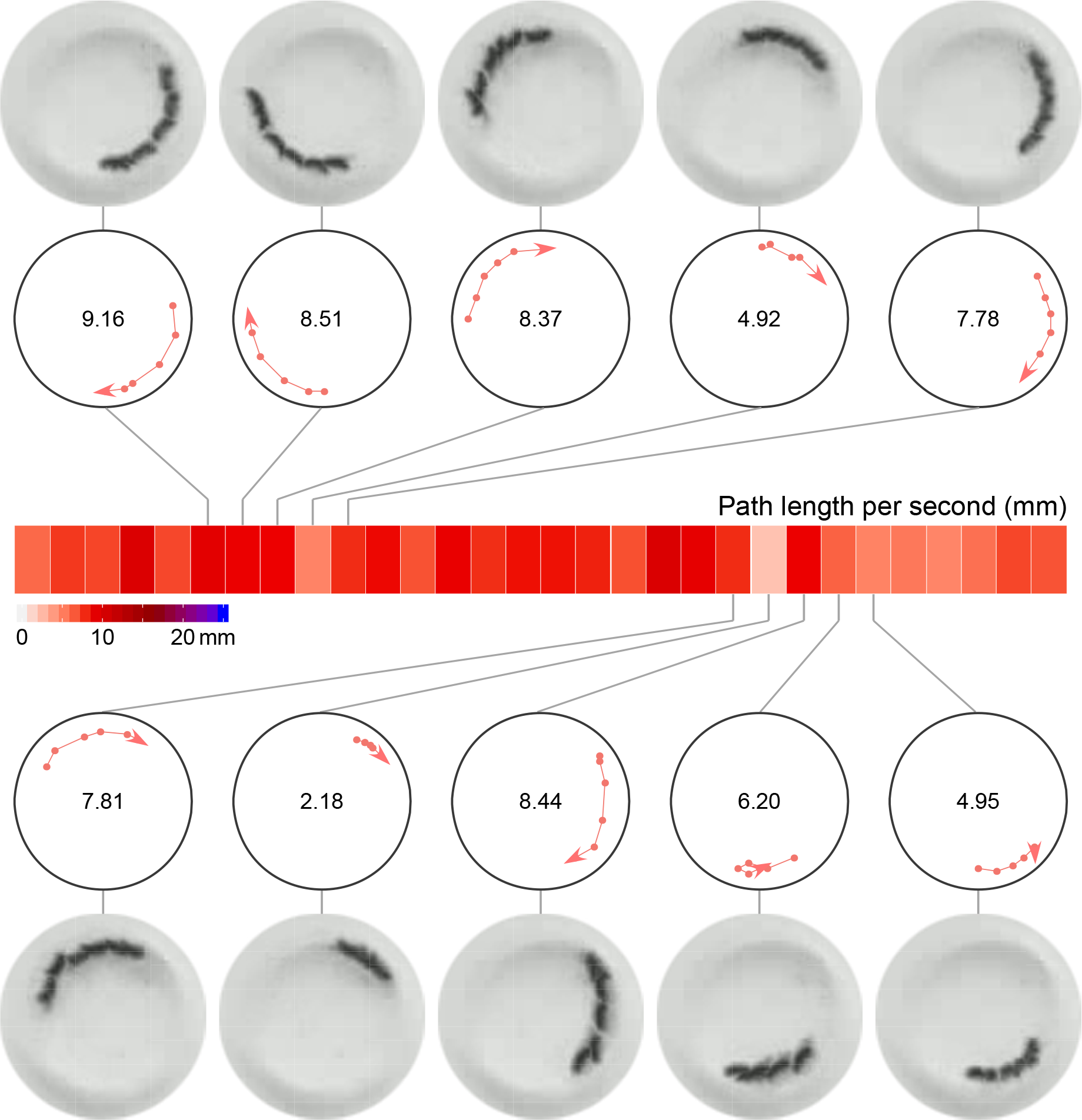
Generation of a heat map. The path length per second (PPS) was calculated based on the fly positions in a circular arena. Fly positions were tracked once every 0.2 s. Each PPS was the sum of travel distances among six consecutive tracking points. Example values pf PPSs were shown. PPSs were displayed as continuous rectangles with graded color representing the values. A relevant color key was provided. For visual clarification, we also presented the actual frames from the video and associated fly positions in the arena (shown as red dots in a circle). Arrows depict walking directions of flies.

### RT-PCR

RT-PCR was performed to examine the transcriptions of *w*^+^ in Canton-S and *w*^1118^ flies. Samples (40-60 male flies per sample) were homogenized with the 7-ml Dounce homogenizer. Total RNA was extracted with TRIzol Reagent (Catalog # 15596026, Invitrogen) by following the manufacturer’s protocol. RNA was then reverse transcribed to cDNA using LunaScript RT SuperMix kit (Catalog # E3010L, New England BioLabs). *w*^1118^ flies might have partial or truncated transcripts of *w*^+^. Therefore, to gain better information we designed three pairs of primers: wF1 (5’-AGTGCTGTGCCAAAACTCCT-3’) and wR1 (5’-ATTGACCGCCCCAAAGATGT-3’); wF2 (5’-CCAAAAACTACGGCACGCTC-3’) and wR2 (5’-GAAAGGCAAGGGCATTCAGC-3’); wF3 (5’-TCTCATGGTTCCGTTACGCC-3’) and wR3 (5’-GCCCGAAGTCTTAGAGCCAG-3’). The transcripts of *Tub84B* were used as a loading control with its own primers: Tub84BF (5’-CATGGTCGACAACGAGGCTA-3’) and Tub84BR (5’-GCACCTTGGCCAAATCACCT-3’). PCR was conducted using the 2×Taq FroggaMix kit (#FBTAQM, FroggaBio) in the S1000 thermal cycler (Catalog # 184-0048, Bio-Rad). The thermal cycling conditions were the following: initial denaturation at 94 °C for 3 min, followed by 35 cycles of denaturation (94 °C 30 s), annealing (58 °C 30 s), and extension (72 °C 60 s), with a final extension at 72 °C for 10 min. The amplified DNA fragments were mixed with Novel juice (Catalog # LD001-1000, FroggaBio) at 5:1, separated in 2 % gel of agarose (A87-500G, FroggaBio), and visualized with AlphaImager 2200 Gel Documentation and Image Analysis System (Alpha Innotech).

### Immunofluorescence

The detailed procedure has been described (Xiao, 2016). Briefly, dissected brain tissues were fixed in 4% paraformaldehyde at room temperature for 1h. After three times of washes with PBT (1×PBS with 0.5 % BSA, 0.5 % Triton X-100), tissues were incubated with a saturation buffer (PBT containing 10 % normal goat serum) for 1h at room temperature. Samples were then incubated with the primary antibody (mouse anti-GFP, catalog# 12A6, DSHB) at 1:20 (diluted with saturation buffer) in the cold room (4 °C) overnight. After three washes, tissues were incubated in the secondary antibody (Alexa Fluor-488 conjugated goat anti-mouse IgG, 115-545-003, Jackson ImmunoResearch) at 1:500 in the cold and darkroom for 12 - 24 h. After washes, tissues were suspended in 200 μl SlowFade Gold Antifade Mountant (S36940, Life Technologies) and mounted on slides for microscopy. Image stacks were taken with a laser scanning microscope (Carl Zeiss LSM 710NLO) and associated software Zen 2009 (Carl Zeiss). Image processing was performed with ImageJ (Schneider et al., 2012).

### Statistics

Heat maps were used for graphic representation of PPSs with color gradients. Density plots were performed for the visualization of PPSs distribution. Mann-Whitney U test was conducted to compare the difference of 5-min path length between two groups. Kruskal-Wallis rank sum test and the post-hoc Dunn test were performed to examine the difference of 5-min path length in more than three groups of flies. Ridgeline plot was carried out to show the distributions of the dataset of 300 PPSs for each genotype. Spearman correlation analysis was used to evaluate the association between *w*^+^/m*w*^+^ copy number and median PPS. Fourier transform was conducted to produce the frequency domain of data from a time series (i.e. PPSs over time). Generalized additive models (GAM) with integrated smoothness estimation was used to generate the smoothed regression lines for power spectrum analysis. The stacked bar plot was used to show the percentages of locomotor components with different frequencies. Post-hoc pairwise Fisher’s exact tests were performed to compare the proportions of high-frequency components among multiple groups. Compact letter display was used to show the difference from all pare-wise comparisons by letters. Statistics are performed with the software R (R Core Team, 2019). *P* < 0.05 is considered statistically significant.

## Results

### *w*^+^ locus suppressed locomotion

We examined the walking activities of individual adult flies in circular arenas (12.7 mm diameter 3 mm depth). A parameter, path length per second (PPS), was calculated and the sequential PPSs during a period of time were shown (Fig. 2A). Wildtype Canton-S males showed the overall PPSs relatively short compared with the males of *w*^1118^, a white-eyed mutant. Canton-S flies walked relatively intermittently (noted as active walking interspaced by inactive or resting episodes) while *w*^1118^ flies walked continuously. *w*^+^ F10 flies (which contained the *w*^+^ locus in the *w*^1118^ isogenic background after ten-generation serial backcrossing) walked intermittently, a pattern observed in Canton-S. This observation indicated an association between *w*^+^ locus and the pattern of intermittent walking.

**Figure 2.**
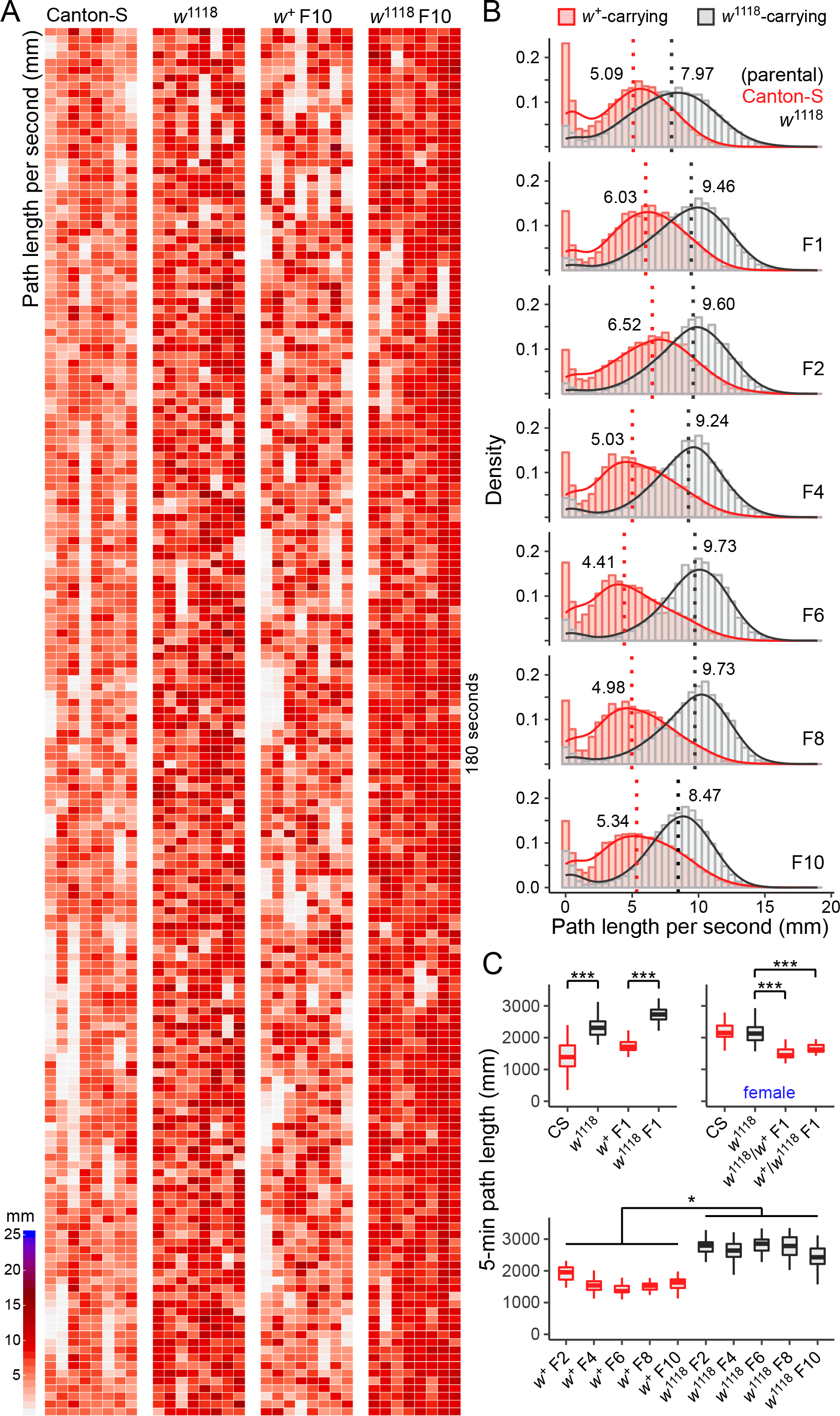
*w*^+^ locus suppressed the locomotion of adult flies in circular arenas. (**A**) Heat maps of path length per second (PPS) from four groups of flies (Canton-S, *w*^1118^, *w*^+^ F10, and *w*^1118^ F10). Data of 180 seconds of activities are presented. (**B**) Density plots of path length per second from *w*^+^-carrying (red) and *w*^1118^-carrying flies (grey). F1-10 flies are the offspring from serial backcrossing. Dotted lines represent median PPSs (values are shown). (**C**) Plots of 5-min path length among different groups of flies (*w*^+^-carrying flies in red bars; *w*^1118^-carrying flies in black bars). Most data are from male flies. Data from female flies are indicated. * *P* < 0.05, *** *P* < 0.001 by Kruskal-Wallis rank sum test and the post-hoc Dunn test.

In contrast, *w*^1118^ F10 flies walked relatively continuously, as *w*^1118^ flies did. *w*^1118^ F10 flies have lost *w*^+^ locus but retained wildtype genetic background compared with Canton-S. The observations in *w*^1118^ F10 confirmed the association between *w*^+^ locus and intermittent walking pattern. Further analysis in F1-8 flies showed that all *w*^+^-carrying males performed intermittent walking whereas all *w*^1118^-carrying males performed relatively continuous walking (Fig. S1).

These results suggested the suppression of locomotion by *w*^+^ locus. More specifically, *w*^+^ locus might be associated with the generation of the intermittent walking pattern.

Density plots of PPSs showed that Canton-S males had a median PPS (5.09 mm) relatively small compared with *w*^1118^ males (7.97 mm) (Fig. 2B). In fact, *w*^+^-carrying males of all tested generations (i.e. F1, F2, F4, F6, F8 and F10) had smaller median PPSs than their *w*^1118^-carrying counterparts. Additionally, *w*^+^-carrying males displayed a secondary peak of PPS near zero, indicating more resting during locomotion than *w*^1118^-carrying males.

Notably, Canton-S females displayed a relatively continuous walking pattern that was similar to *w*^1118^ females and their median PPSs were also comparable (Canton-S 7.53 mm, *w*^1118^ 7.25 mm) (Fig. S2). *w*^+^ F10 and *w*^1118^ F10 females, however, showed intermittent and continuous walking patterns, respectively. *w*^+^ F10 females had a small median PPS (5.46 mm) relative to *w*^1118^ F10 females (8.98 mm). Both female F1 hybrids (*w*^1118^/*w*^+^ F1 and *w*^+^/*w*^1118^ F1, the former in the wildtype cytoplasmic background the latter *w*^1118^ cytoplasmic background) that carried the heterozygous *w*^+^ allele displayed similar patterns of intermittent walking. In general, these results firmly supported the suppression effect of *w*^+^ locus on locomotion. The only exception, the continuous walking pattern in Canton-S females, might be a consequence of sexual dimorphism or long-term laboratory adaptation.

The 5-min path lengths were compared. Canton-S males had shorter 5-min path lengths than *w*^1118^ males (*P* < 0.001) (Fig. 2C). *w*^+^ F1 males had shorter path lengths than *w*^1118^ F1 males (*P* < 0.001). In females, there was no statistical difference of 5-min path length between Canton-S and *w*^1118^. This was different from the observations of males. There was also an insignificant difference in 5-min path length between the female F1 hybrids. Both hybrids carried heterozygous *w*^+^ allele and showed shorter 5-min path lengths than *w*^1118^ flies (*P* < 0.001 for both comparisons), supporting the notion that *w*^+^ locus (but not the cytoplasmic background) had a suppression effect on the locomotion. *w*^+^ F10 females, however, displayed shorter 5-min path lengths than *w*^1118^ F10 females (*P* < 0.001) (Fig. S2). The suppression effect was further observed that all other tested *w*^+^-carrying males (i.e. F2, F4, F6, F8 and F10) had shorter 5-min path lengths than *w*^1118^-carrying males (*P* < 0.05).

### Reduced transcription of *w*^+^ in *w*^1118^ flies

Three pairs of primers were used to examine the transcription of *w*^+^ in Canton-S and *w*^1118^ flies. Compared with Canton-S, transcription of *w*^+^ in *w*^1118^ flies was reduced mainly in the first (5’) half of the gene while the transcription of the last exon was likely unchanged (Fig. 3). These results were consistent with previous reports (Dalby and O’Hare, 1991; Rabinow and Birchler, 1989).

**Figure 3.**
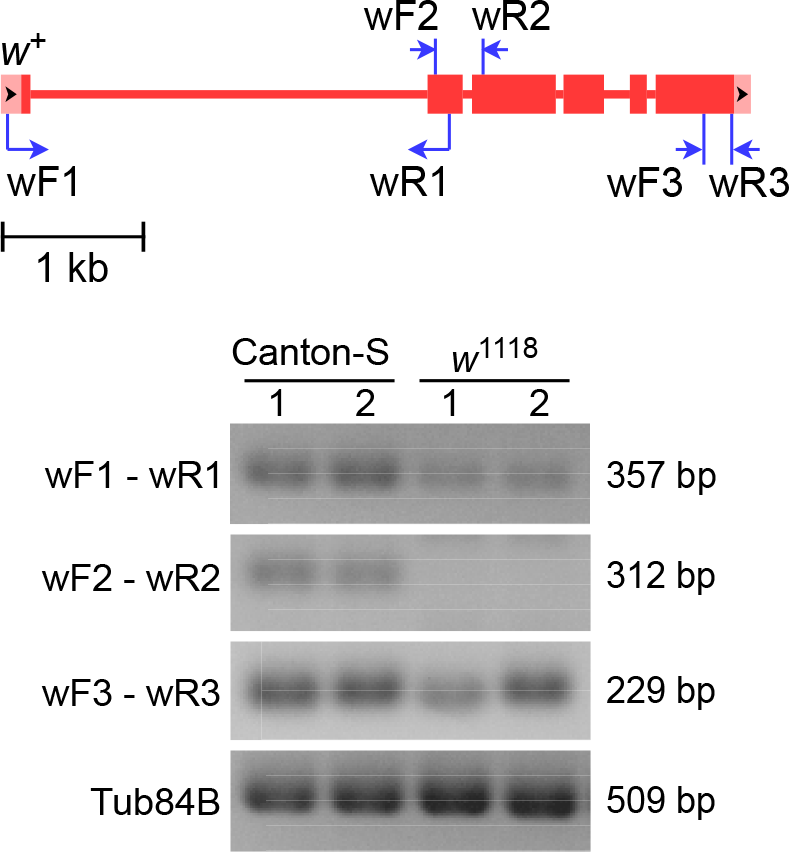
Reduced transcription of the *w*^+^ gene in *w*^1118^ flies. Top panel: The structure of *w*^+^ and primer-specific locations. Bottom panel: Gel images of RT-PCR products by three sets of primers for *w*^+^ and by a pair of primers for the loading control *Tub84B*. Sizes of amplified DNA products are provided.

### m*w*^+^ suppressed locomotion in a manner of copy-number-dependent

We next examined whether the marker gene m*w*^+^, a miniature form of *w*^+^, was associated with the locomotor suppression. The UAS flies containing 1-4 genomic copies of m*w*^+^ at fixed recombination sites (i.e. attP2 and attP40) were produced. Before recombination, all the UAS flies have been backcrossed into *w*^1118^ flies for ten generations (Xiao and Robertson, 2016).

Flies carrying one genomic copy of m*w*^+^ displayed slightly reduced median PPSs compared with *w*^1118^ flies (Fig. 4A). Flies with increased genomic copies of m*w*^+^ had gradually decreased median PPSs. There was a negative correlation between the copy number of m*w*^+^ and median PPS (*r* = −0.94, *P* < 0.0001) (Fig. 4B). Interestingly, flies with a single genomic copy of *w*^+^ that was duplicated to the Y chromosome had median PPSs comparable to those with one genomic copy of m*w*^+^. Therefore, one copy of m*w*^+^ and the duplicated *w*^+^ likely had similar effects on the locomotion. Flies containing an increased copy number of m*w*^+^ displayed increased chances of intermittent walking (Fig. S3).

**Figure 4.**
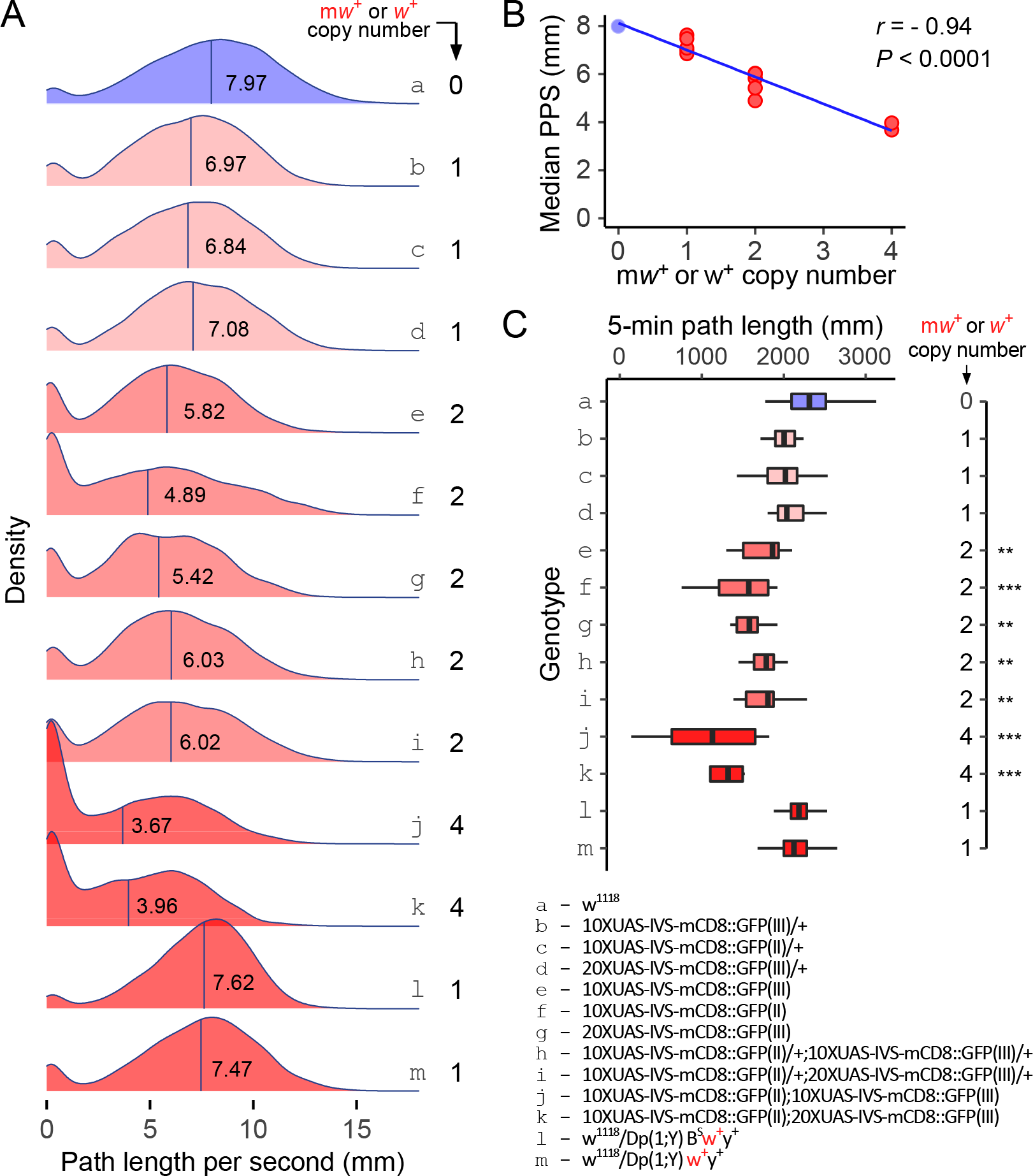
Multiple genomic copies of m*w*^+^ suppressed the locomotion of adult flies. (**A**) Density ridges of path length per second in flies of different genotypes. The genomic copy number of m*w*^+^ or *w*^+^ is provided. Color gradients represent roughly the copy number of m*w*^+^ or *w*^+^. *w*^1118^ flies (with blue color under the ridge) have no m*w*^+^ (indicated as 0 copy). Lower-case letters represent different genotypes which are indicated in (**C**). (**B**) Analysis of correlation between the copy number of m*w*^+^ (or duplicated *w*^+^) and median PPS. Spearman correlation analysis was used. (**C**) Comparison of 5-min path length among flies with different genotypes. The copy number of m*w*^+^ or *w*^+^ for each genotype is shown. Lower case letters represent different genotypes. ** *P* < 0.01, *** *P* < 0.001 by Kruskal-Wallis rank sum test and the post-hoc Dunn test.

Flies carrying a single copy of m*w*^+^ or Y-duplicated *w*^+^ had the same levels of 5-min path length as *w*^1118^ flies (Fig. 4C). Flies with two to four copies of m*w*^+^ had reduced 5-min path lengths compared with *w*^1118^ flies. Therefore, multiple copies of m*w*^+^ were associated with shorter median PPS and reduced 5-min path length. These data suggested that m*w*^+^ suppressed locomotion in a manner of copy-number-dependent.

### *w*^+^ selectively suppressed high-frequency components of locomotion

The dataset of sequential PPSs over time could be regarded as a time series that can be decomposed by Fourier transform into a series of frequency components. Each frequency and its associated power strength are representative of a structural unit. Here, we treated such a structural unit as a “locomotor component” since it contained information that could sufficiently describe a subset component of behavior from the complicated locomotor activity. *w*^+^ could suppress the locomotor components selectively.

A full-spectrum analysis of the time-series (i.e. PPSs over 300 s) was performed (Fig. 5A, Fig. S4). Common to all tested flies, locomotor components at relatively low frequencies (0.0067-0.1 Hz) had high power whereas components at high frequencies (0.1-0.5 Hz) had low power. Canton-S showed a great reduction of the power spectrum at nearly all frequencies compared with *w*^1118^ flies. *w*^+^ F10 flies showed reduced power spectrum at high frequencies and increased power at low frequencies compared with *w*^1118^ flies. Female flies of Canton-S, *w*^1118^, *w*^+^ F10 displayed the power-frequency relations that were similar to males (Fig. S5). *w*^+^ F10 flies differed from *w*^1118^ that the former carried a *w*^+^ locus. Therefore, *w*^+^ locus was associated with a decreased power spectrum at relatively high frequencies and an increased power spectrum at low frequencies.

**Figure 5.**
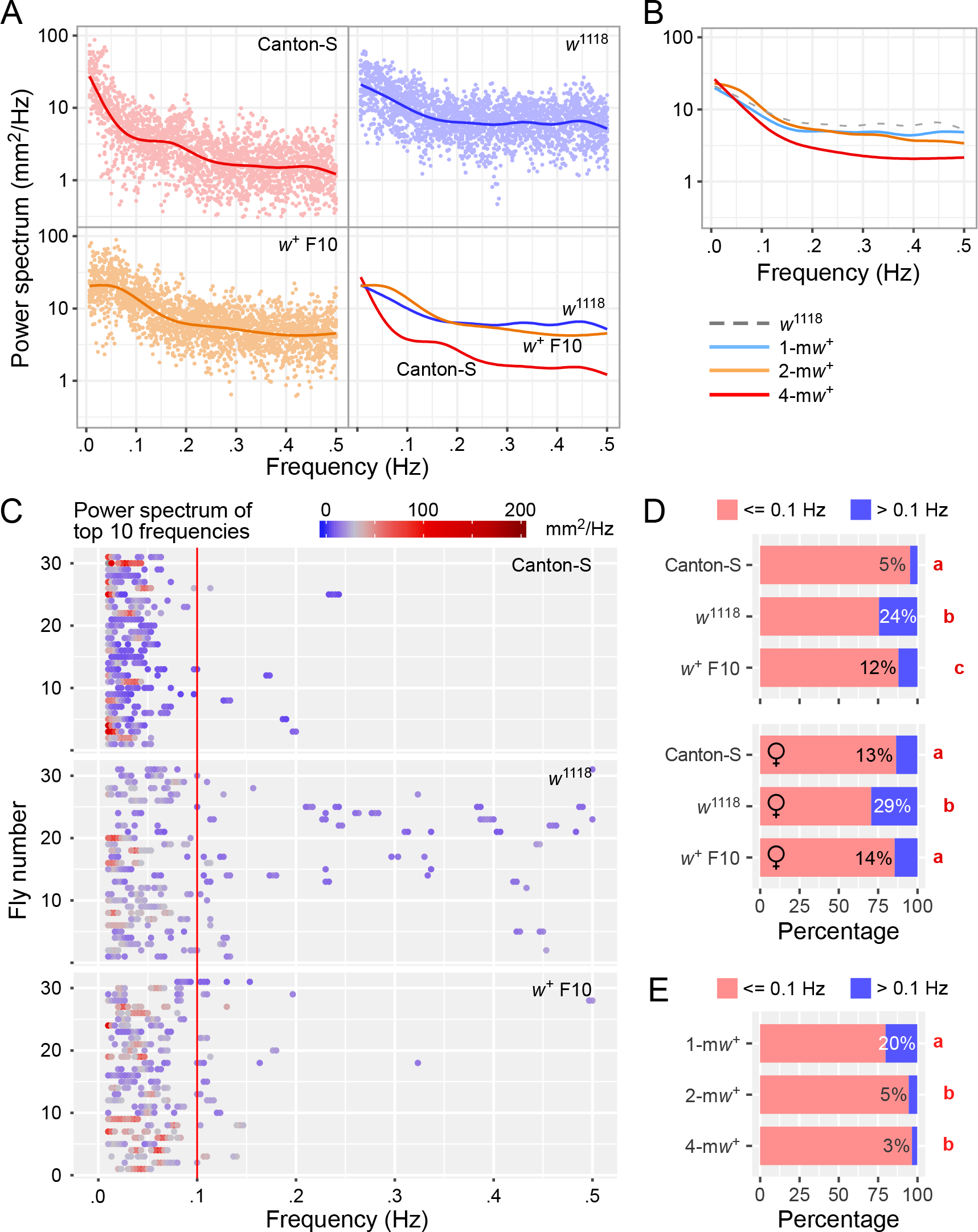
*w*^+^ selectively suppressed locomotor components at relatively high frequencies. (**A**) Power spectrum analysis of locomotor components at different frequencies in male flies. The frequency-domain data (with a full spectrum at 0.0067-0.5 Hz) are generated by Fourier transform. The scatter plot and a smoothed regression line are provided for each genotype. Only smoothed regression lines are shown for the overlapping plot of three genotypes (Canton-S, *w*^1118^, *w*^+^ F10). The smoothed regression lines were generated by fitting the generalized additive model to data. (**B**) Power spectrum analysis of locomotor components in m*w*^+^-carrying flies. Flies carrying one genomic copy of m*w*^+^ (1-m*w*^+^): 10×UAS -IVS-mCD8∷GFP (attP2)/+, 10×UAS -IVS-mCD8∷GFP (attP40)/+, and 20×UAS -IVS-mCD8∷GFP (attP2)/+. Flies carrying two genomic copies of m*w*^+^ (2-m*w*^+^): 10×UAS -IVS-mCD8∷GFP (attP2), 10×UAS -IVS-mCD8∷GFP (attP40), 20×UAS -IVS-mCD8∷GFP (attP2), 10×UAS -IVS-mCD8∷GFP (attP40)/+; 10×UAS -IVS-mCD8∷GFP (attP2)/+, and 10×UAS -IVS-mCD8∷GFP (attP40)/+; 20×UAS -IVS-mCD8∷GFP (attP2)/+. Flies carrying four genomic copies of m*w*^+^ (4-m*w*^+^): 10×UAS -IVS-mCD8∷GFP (attP40); 10×UAS -IVS-mCD8∷GFP (attP2), and 10×UAS -IVS-mCD8∷GFP (attP40); 20×UAS -IVS-mCD8∷GFP (attP2). The dotted line (data of *w*^1118^) is duplicated from (A) for statistical convenience. (**C**) Power spectrum analysis of top-ten frequencies. The top-ten frequencies were sampled from locomotor components that carried ten highest power spectrums. Color gradients indicate the power strength for a specific frequency. A color key is provided. For each genotype, data from 31 flies are shown. Genotypes are indicated. (**D**) Stack-bar plots of percentages of locomotor components at different frequencies. The provided values are the percentages of high-frequency components. Low frequency range: <= 0.1 Hz (red); High frequency range: > 0.1 Hz (blue). Fly genotypes are indicated. Lower-case letters indicate statistical significance at *P* < 0.05 (from an R function of compact letter display). (**E**) Stack-bar plots of percentages of locomotor components at different frequencies in m*w*^+^-carrying flies. See (B) for the detailed genotypes for 1-m*w*^+^, 2-m*w*^+^, and 4-m*w*^+^ flies.

The 1-m*w*^+^ flies (which carried one genomic copy of m*w*^+^) showed a slightly decreased power spectrum at high frequencies. The 2-m*w*^+^ flies showed further reduction of the power spectrum at high frequencies and an increase at relatively low frequencies. The 4-m*w*^+^ flies displayed a greater reduction of the power spectrum at high frequencies (Fig. 5B). Data suggested that *w*^+^ had a suppression effect on the high-frequency components of locomotion.

We quantified the frequency distribution by sampling the top-ten components that carried ten highest power spectrums. The frequency distribution of top-ten components could be reflective of the composition of major locomotor structures. Most of the top-ten components were at a range of low frequencies (0.0067-0.1 Hz) in the tested flies (Fig. 5C). *w*^1118^ flies showed clearly an increase of components at high frequencies compared with Canton-S. *w*^+^ F10 flies showed a reduction of high-frequency components compared with *w*^1118^ flies. Statistically, *w*^1118^ flies displayed 24% of locomotor components at high frequencies (> 0.1 Hz), which was higher than Canton-S (5%) (*P* < 0.05, Post-hoc pairwise Fisher’s exact tests) and also higher than *w*^+^ F10 (12%) (*P* < 0.05) (Fig. 5D). The reduction of the power spectrum at relatively high frequencies in *w*^+^-carrying flies was also observed in female flies. Results indicated that *w*^+^ locus selectively suppressed the high-frequency components.

The 1-m*w*^+^ flies had 20% of top-ten components at high frequencies, which was statistically the same as *w*^1118^ (24%) (*P* > 0.05). The 2-m*w*^+^ flies showed reduced locomotor components at high frequencies (5%) and 4-m*w*^+^ flies showed even greater reduction (3%) compared with *w*^1118^ (Fig. 5E). Thus, like *w*^+^ locus, multiple copies of m*w*^+^ selectively suppressed high-frequency components of locomotion.

### *w*^+^ knockdown in neurons increased the proportion of high-frequency locomotor components

We next examined the effects of *w*^+^ downregulation on the frequency distribution of top-ten locomotor components. *w*^+^ knockdown in neurons (by the driver elav-Gal4) resulted in greatly reduced power at low frequencies relative to the controls (Fig. 6A). In contrast, the elav-Gal4-driven expression of a membrane protein mCD8-GFP had little effect on the power spectrum at low frequencies (Fig. 6B). Downregulation of *w*^+^ in glial cells (by repo-Gal4) resulted in no reduction of the power spectrum at low frequencies (Fig. 6C). Notably, since both GAL4/UAS flies and Gal4 controls carried 2-3 genomic copies of m*w*^+^, the probable confounding effects (i.e. suppression of high-frequency components by multiple m*w*^+^) were appropriately balanced between the experimental and control flies.

**Figure 6.**
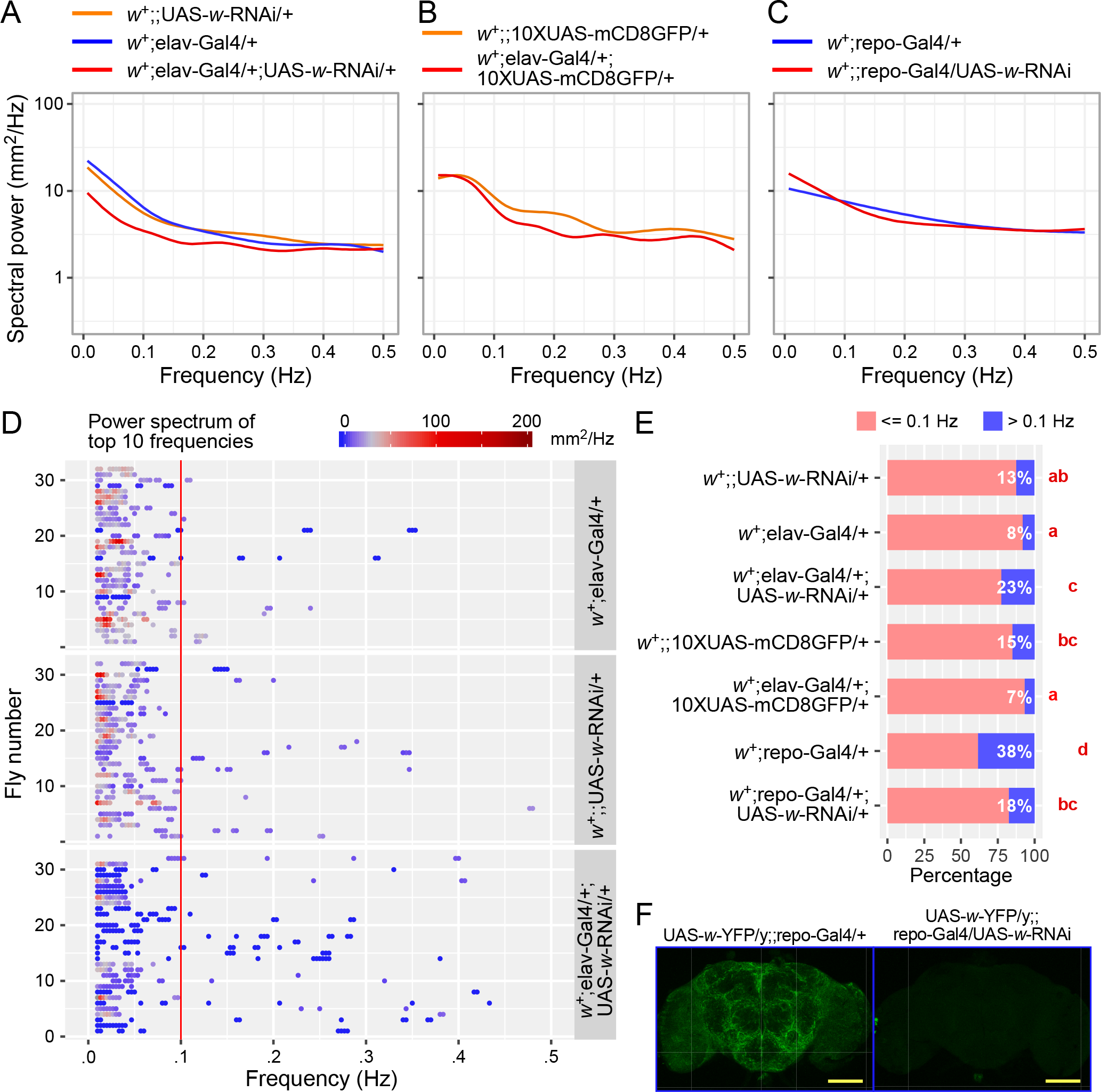
*w*^+^ knockdown in neurons resulted in an increased proportion of high-frequency locomotor components. (**A**) RNAi knockdown of *w*^+^ in neurons (*w*^+^; elav-Gal4/+; UAS-*w*-RNAi/+) caused a reduction of locomotor components mainly at relatively low frequencies (<= 0.1 Hz). (**B**) Expression of an ectopic membrane protein mCD8-GFP by the same Gal4-driven expression system caused no effect on locomotion. (**C**) RNAi knockdown of *w*^+^ in glial cells *w*^+^;;repo-Gal4/UAS-*w*-RNAi) caused no effect on locomotion. (**D**) Power spectrum analysis of top-ten frequencies. RNAi knockdown of *w*^+^ in neurons (*w*^+^; elav-Gal4/+; UAS-*w*-RNAi/+) caused increased components at high frequencies (> 0.1 Hz). Color gradients indicate the power strength for a specific frequency. A color key is provided. For each genotype, data from 31 flies are shown. (**E**) Stack-bar plots of percentages of locomotor components at different frequencies. Low frequency range: <= 0.1 Hz (red); High frequency range: > 0.1 Hz (blue). The provided values are the percentages of high-frequency components. Lower-case letters indicate statistical significance at *P* < 0.05 (from an R function of compact letter display). (**F**) Glial expression of the White protein (UAS-*w*-YFP/y;;repo-Gal4/+) was inhibited by the *w*-RNAi (in UAS-*w*-YFP/y;;repo-Gal4/ UAS-*w*-RNAi). Image data from a PhD thesis (Xiao, 2016) with permission. Scale bar 100 μm.

Flies with *w*^+^ knockdown in neurons (*w*^+^; elav-Gal4/+; UAS-*w*-RNAi/+) displayed an increased proportion of top-ten components at high frequencies (Fig. 6D and 6E). Neither the expression of mCD8-GFP in neurons nor downregulation of *w*^+^ in glia had such an effect. The *w*^+^ knockdown was evident by the observation that the glial expression of a recombinant *w* was blocked by the RNAi specific to this gene (Fig. 6F).

## Discussion

We report that the *w*^+^ gene modulates the walking behavior of adult flies by selectively suppressing high-frequency locomotor components. We initially observed two different locomotor patterns in adult flies having different *w* alleles: intermittent walking in *w*^+^-carrying flies and continuous walking in *w*^1118^-carrying flies. We then show that both *w*^+^ locus and multiple genomic copies of m*w*^+^ reduce median PPSs and 5-min path length and importantly, the suppression effect of m*w*^+^ is dependent on the copy number. The Fourier transform of time series (PPSs over time) reveals that *w*^+^ re-shapes the frequency distribution of locomotor structures by specifically suppressing components at relatively high frequencies (i.e. 0.1-0.5 Hz). Lastly, the downregulation of *w*^+^ in neurons causes an increased proportion of high-frequency locomotor components.

Extensive effort has been made to control the genetic or cytoplasmic background in order to better understand the behavioral consequences of the *w*^+^ gene (Xiao and Robertson, 2015; Xiao and Robertson, 2016). Different from the pattern of continuous walking in *w*^1118^ flies, the intermittent walking in Canton-S flies could be associated with any of the three main factors: the *w*^+^ allele, its wildtype genetic background, and wildtype cytoplasmic background. Since *w*^1118^ flies contain a mutant *w* allele, an isogenic genetic background, and probably a different cytoplasmic background, a direct comparison between *w*^1118^ and Canton-S is inappropriate. Perhaps the best-controlled comparison is between the pair of *w*^+^ F10 and *w*^1118^. After ten-generation backcrossing, the established *w*^+^ F10 flies have included the *w*^+^ locus into the isogenic genetic background while the cytoplasmic background remains the same to *w*^1118^ flies. *w*^+^ F10 flies have retained the pattern of intermittent walking but not the continuous walking. Therefore, there is a strong association between the pattern of intermittent walking and *w*^+^ locus.

Further observations have supported this association. From F1 to F10, *w*^+^-carrying flies have their mixed genetic background (half wildtype half isogenic) gradually replaced with the isogenic background. During the process of serial backcrossing, the association between *w*^+^ locus and intermittent walking is persistently retained. Similarly, the association between *w*^1118^ locus and continuous walking is sustained. The tight relationship between *w*^+^ locus and intermittent walking can also be found by the comparison of two types of female F1 hybrids. These two types of females have the same genetic content (heterozygous *w* alleles and heterozygous genetic background) but different cytoplasmic background. Both F1 hybrids, however, display intermittent walking patterns. These data rule out the contribution of cytoplasmic background and confirm that *w*^+^ locus is responsible for the intermittent walking.

The tested m*w*^+^-carrying flies are the UAS lines with fixed docking sites (either attP2 or attP40) where the m*w*^+^-bearing DNA constructs are inserted. Therefore, the potential position effects of m*w*^+^ expression are well-controlled. In addition, UAS flies have been synchronized into *w*^1118^ isogenic background for at least ten generations before the chromosomal recombination (for producing flies with multiple genomic copies of m*w*^+^). Moreover, in the absence of the Gal4 transcriptional factor, the transgenes in the UAS constructs remain silent and thus the confounding factors have been greatly avoided or reduced. After such intensive genetic manipulations, we observe the intermittent walking in flies carrying multiple genomic copies of m*w*^+^ and the suppression of m*w*^+^ on the locomotor parameters. Most interestingly, the suppression effect is dependent on the copy number of m*w*^+^ in the genome. Therefore, the suppression effect on locomotion is specifically associated with the *w*^+^ gene. Notably, the effects of duplicated *w*^+^ on the locomotor parameters (i.e. median PPS and 5-min path length) mimic those of a single genomic copy of m*w*^+^, suggesting a similar function between m*w*^+^ and translocated *w*^+^. Together, these findings indicate that the *w*^+^ gene itself has a suppression effect on locomotion.

The observable exemption exists in Canton-S females which display continuous walking, a characteristic pattern found in both males and females of *w*^1118^ and all tested homozygous *w*^1118^-containing F1-10 males. It is possible that there is a sexual dimorphism of locomotor pattern in wildtype, but this sexual dimorphism might have been lost or modified in *w*^1118^ flies. Alternatively, the long-term laboratory maintenance of fly stocks may have posed an adaptive behavioral response that could be different between fly strains.

Analysis of the frequency domain of locomotion has revealed that common to all tested flies, locomotor components with high power have converged to a narrow range of relatively low frequencies at 0.0067-0.1 Hz. In other words, major behavioral components of walking activities appear to be rhythmic with the periodicities of 10-150 s. Canton-S flies have highly converged frequencies for the top-ten locomotor components thus the locomotion (reflected as sequential PPSs) would appear to be highly rhythmic. This explains the observed intermittent walking of Canton-S males in the circular arenas. In contrast, *w*^1118^ flies have relatively broad frequency distribution of top-ten locomotor components and thus their sequential PPSs could be non-rhythmic and smearing-like. This is consistent with the observation of continuous walking patterns in *w*^1118^-carrying flies. Apparently, a shift of locomotor pattern from continuous walking (found in *w*^1118^-carrying flies) to intermittent walking (found in *w*^+^-carrying flies) would require the suppression of locomotor components at relatively high frequencies (i.e. > 0.1 Hz). Through hybridization, serial backcrossing, and relevant genetic manipulations between Canton-S and *w*^1118^ flies, we show that *w*^+^ locus but not the genetic background or cytoplasmic background is responsible for the suppression of high-frequency locomotor components. Two additional sets of experiments, (1) multiple copies of m*w*^+^ reduce high-frequency locomotor components and (2) downregulation of *w*^+^ in neurons causes an increase of high-frequency components, have provided data to refine our observations that it is the *w*^+^/m*w*^+^ gene that is responsible for the suppression of high-frequency locomotor components.

When individually restrained, Canton-S flies display episodic motor activities with a periodicity of around 19 s (Qiu et al., 2016). The periodicity of episodic motor activity falls within the range of the periodicities of top-ten locomotor components in Canton-S flies. Although different experimental settings are used (episodic motor activities are recorded from the brains of rigidly restrained flies whereas locomotor components are computed from the walking activities of free-moving animals), the high coincidence of periodicities from two types of experiments indicates an intrinsically rhythmic motor ability in wildtype flies.

How does the *w*^+^ gene suppress high-frequency locomotor components? In *Drosophila*, the White protein is an ATP-binding cassette transporter and is a key member of the uptake system. Many small molecules, including pigment precursors, biogenic amines, and metabolic intermediates can be translocated by this system (Anaka et al., 2008; Dreesen et al., 1988; Evans et al., 2008; O’Hare et al., 1984; Sullivan and Sullivan, 1975; Sullivan et al., 1979). It has been shown that White interacts with a second messenger, the cyclic guanosine monophosphate (cGMP), in the regulation of recovery time from anoxia in adult flies (Xiao and Robertson, 2017). cGMP modulates the avoidance behavior to hypoxia by manipulating the cGMP-gated cation channel A in *Drosophila* larvae (Vermehren-Schmaedick et al., 2010). These studies provide clues that the White protein might regulate the rates of cGMP release and suppress the spontaneous cGMP actions. Future work should address how White modifies the neural functions that produce rhythmic walking locomotion in adult flies.

## Acknowledgments

The authors are grateful to Dr. Christopher D. Moyes for providing the lab facilities for RT-PCR experiments, to Dr. R Meldrum Robertson for funding support (the Natural Sciences and Engineering Research Council of Canada, RGPIN 40930-09), to the Department of Biology at Queen’s University for the support of confocal imaging, to the Bloomington *Drosophila* Stock Center for providing fly stocks, and to the Developmental Studies Hybridoma Bank (DSHB) for providing the primary antibody.

## Contributions

C.X. conceived the experiments; C.X. and S.Q. conducted the experiments, analyzed the data, and wrote the manuscript. Both authors reviewed the manuscript.

---

**Figure S1.**
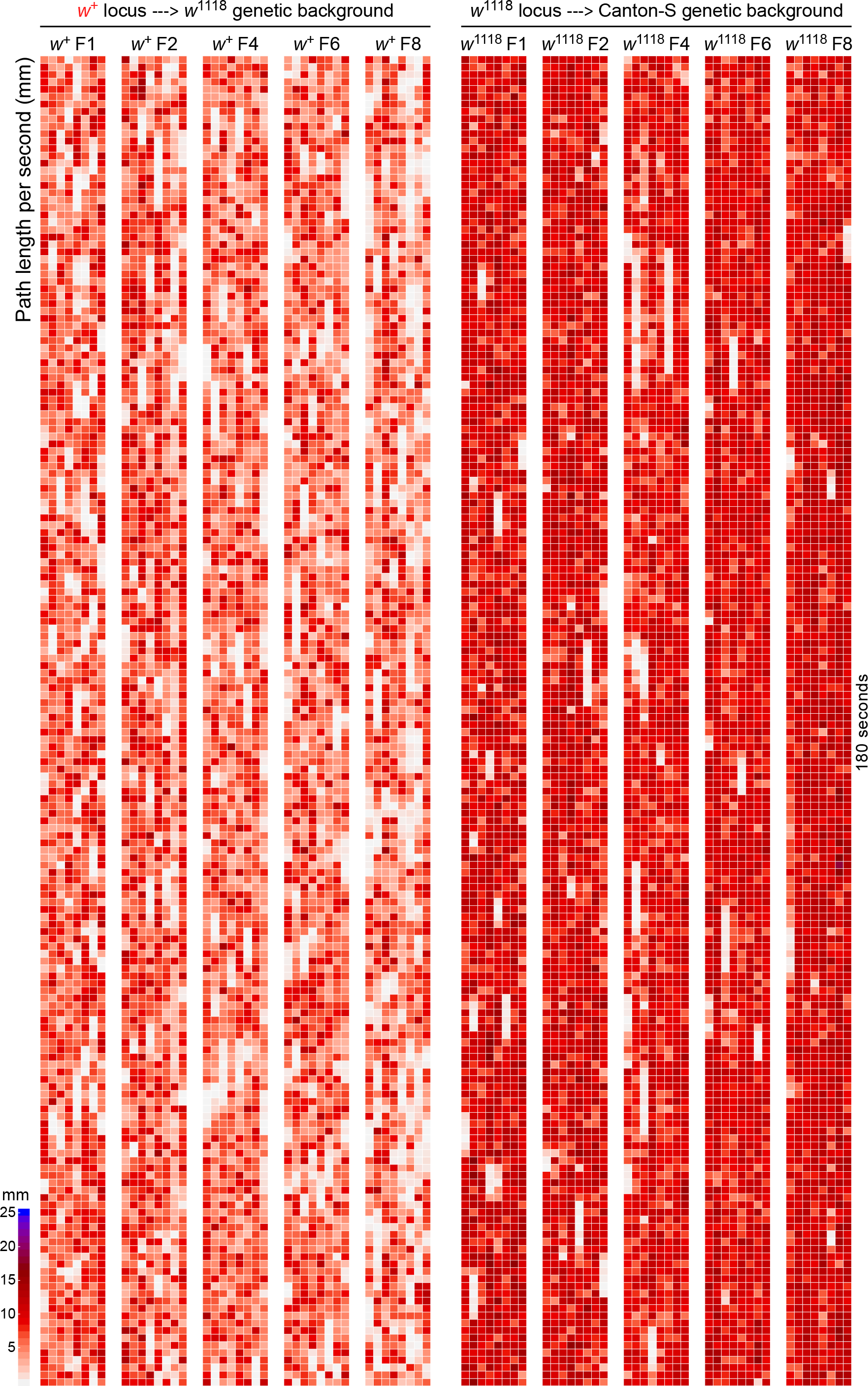
Heat maps of path length per second in flies generated by serial backcrossing. Two different processes of serial backcrossing are shown on the top. F1, F2, F4, F6, and F8 indicate the generations during the backcrossing. Presented data are from male flies. A color key is provided.

**Figure S2.**
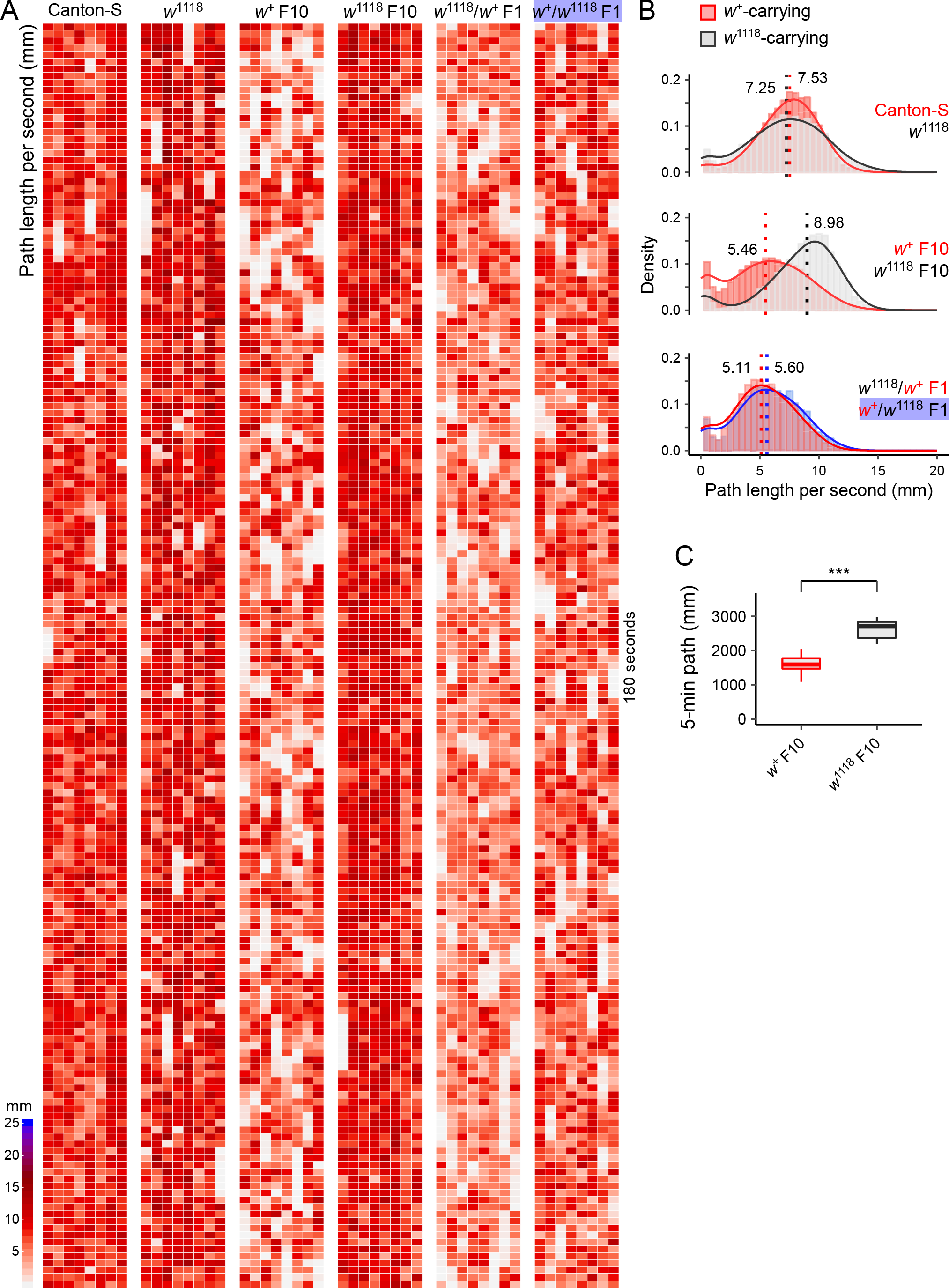
Path length per second in female flies. (**A**) Heat maps of path length per second (PPS) from six groups of female flies (Canton-S, *w*^1118^, *w*^+^ F10, *w*^1118^ F10, *w*^1118^/*w*^+^ F1, and *w*^+^/*w*^1118^ F1). Data of 180 seconds of activities are presented. A color key is provided. The *w*^+^/*w*^1118^ F1 females (blue-shaded) carry the cytoplasmic background from *w*^1118^ flies. (**B**) Density plots of path length per second from female flies. First two plots: *w*^+^-carrying females (red) and *w*^1118^-carrying females (grey); Third plot: F1 hybrid females - *w*^1118^/*w*^+^ F1 contains wildtype cytoplasmic background whereas *w*^+^/*w*^1118^ F1 contains *w*^1118^ cytoplasmic background (blue-shaded). (C) Plots of 5-min path length between the females of *w*^+^ F10 (red) and *w*^1118^ F10 (black). *** *P* < 0.001 by Mann-Whitney U test.

**Figure S3.**
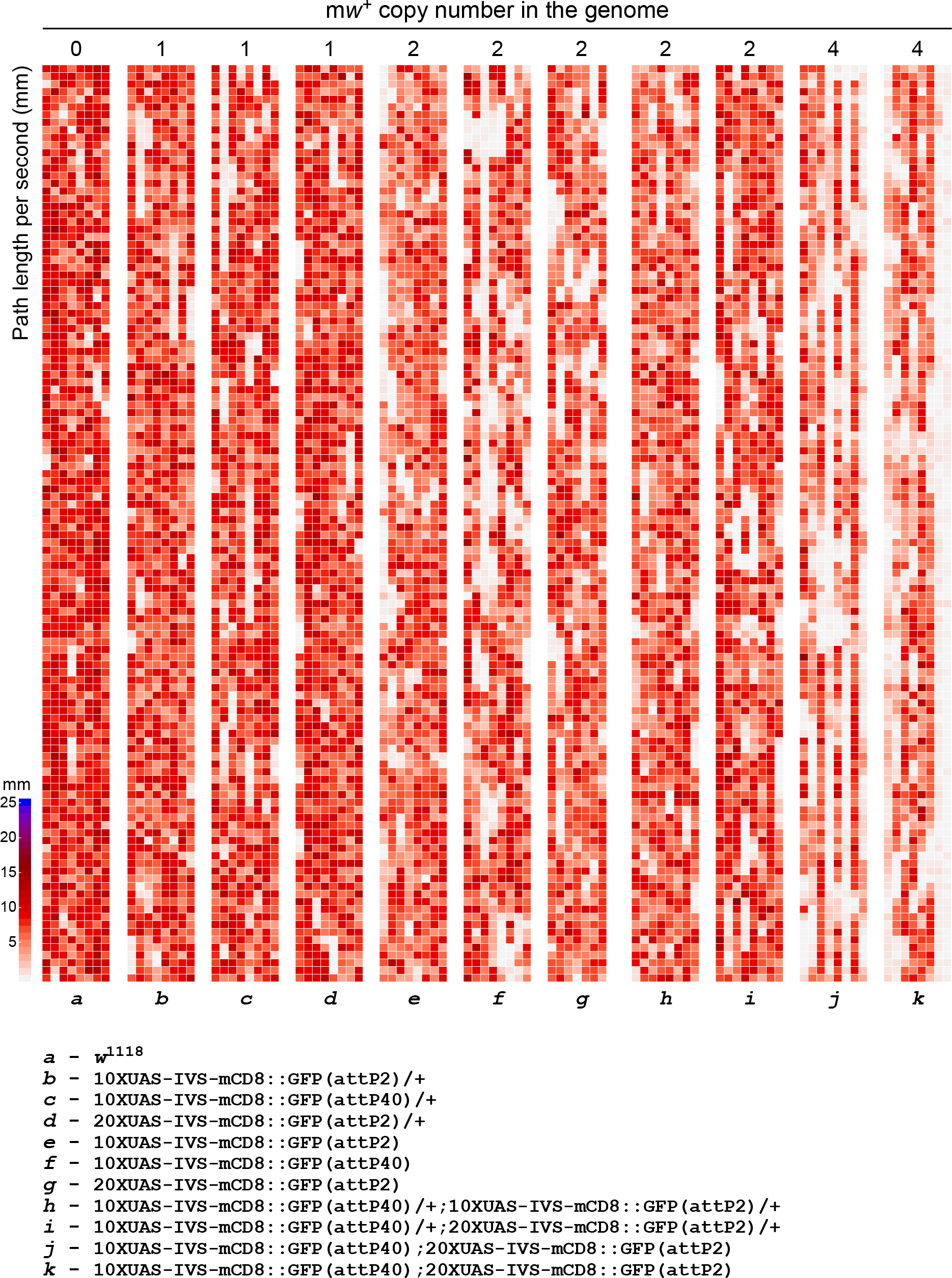
Heat maps of path length per second in m*w*^+^-carrying flies. The m*w*^+^ copy number in the genome are shown on the top. The number 0 indicates *w*^1118^ flies. Lower-cased letters at the bottom are the representations of fly lines whose detailed genotypes are provided. A color key is presented.

**Figure S4.**
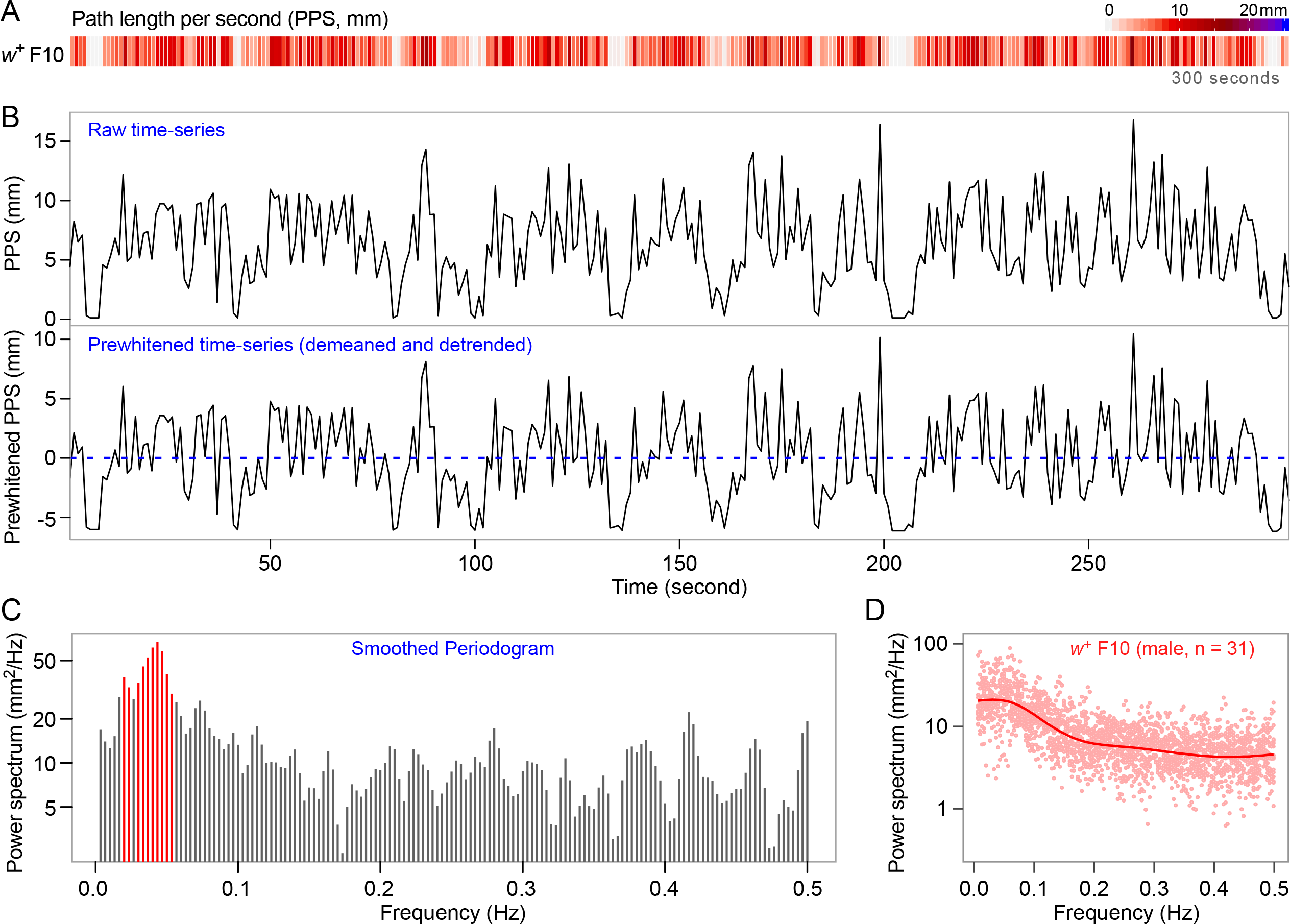
An example of data processing by Fourier transform. (**A**) The heat map of path length per second (over 300 s) from a single male fly of *w*^+^ F10. A color key is provided. (**B**) Plots of path length per second over time. The top plot shows the data of the raw time-series. The bottom plot shows the pre-whitened time-series with the mean and slope adjusted to zero (demeaned and detrended). (**C**) Smoothed periodogram of frequency-domain data which are produced by Fourier transform. Red bars indicate the top-ten frequencies with ten highest power. (**D**) A demonstration of the power spectrum plot and the smoothed regression line from *w*^+^ F10 flies (n = 31). The Fourier transform and data visualization are performed with R.

**Figure S5.**
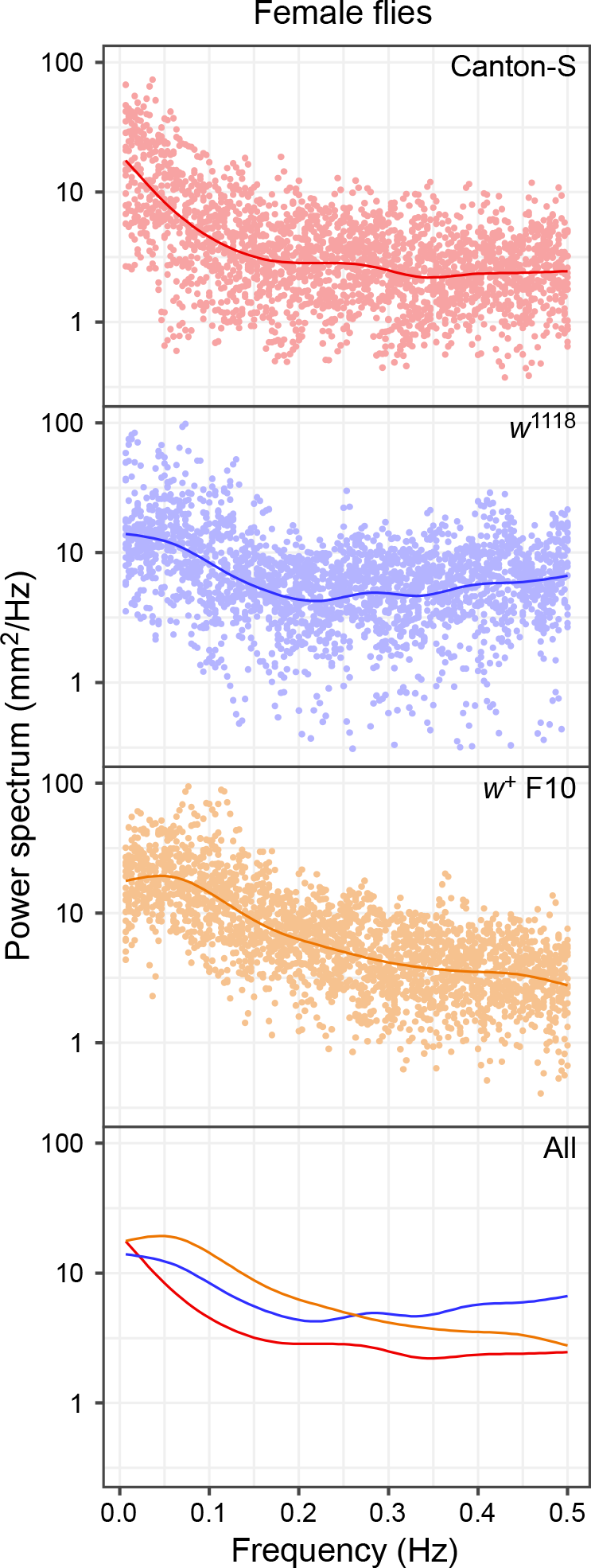
Power spectrum analysis of locomotor components at different frequencies in female flies. The scatter plot and a smoothed regression line are provided for each genotype. Only smoothed regression lines are shown for the overlapping plot of three genotypes (Canton-S, *w*^1118^, *w*^+^ F10). Different colors are used for data visualization.

